# Multimodal layer modelling reveals *in-vivo* pathology in ALS

**DOI:** 10.1101/2023.06.28.546951

**Authors:** Alicia Northall, Juliane Doehler, Miriam Weber, Igor Tellez, Susanne Petri, Johannes Prudlo, Stefan Vielhaber, Stefanie Schreiber, Esther Kuehn

## Abstract

Amyotrophic lateral sclerosis (ALS) is a rapidly progressing neurodegenerative disease characterised by the loss of motor control. Current understanding of ALS pathology is largely based on *post-mortem* investigations at advanced disease stages. A systematic *in-vivo* description of the microstructural changes that characterise early-stage ALS, and their subsequent development, is so far lacking.

Recent advances in ultra-high field (7T) MRI data modelling allow us to investigate cortical layers *in-vivo*. Given the layer-specific and topographic signature of pathology in ALS, we combined submillimeter structural 7T-MRI data (qT1, QSM), functional localisers of body parts (upper limb, lower limb, face) and automated layer modelling to systematically describe pathology in the primary motor cortex (M1), in 12 living ALS-patients with reference to 12 age-, gender-, handedness- and education-matched controls. Longitudinal sampling was performed for a subset of patients. We calculated multimodal pathology maps for each layer (superficial layer, layer 5a, layer 5b, layer 6) of M1 to identify hotspots of demyelination, iron and calcium accumulation in different cortical fields.

We show preserved mean cortical thickness and layer architectures of M1, despite significantly increased iron in layer 6 and significantly increased calcium in layer 5a and superficial layer, in patients compared to controls. The behaviorally first-affected cortical field shows significantly increased iron in L6 compared to other fields, while calcium accumulation is atopographic and significantly increased in the low-myelin borders between cortical fields compared to the fields themselves. A subset of patients with longitudinal data shows that the low-myelin borders are particularly disrupted, and that calcium hotspots but to a lesser extent iron hotspots precede demyelination. Finally, we highlight that a very-slow progressing patient (P4) shows a distinct pathology profile compared to the other patients.

Our data shows that layer-specific markers of *in-vivo* pathology can be identified in ALS-patients with a single 7T-MRI measurement after first diagnosis, and that such data provide critical insights into the individual disease state. Our data highlight the non-topographic architecture of ALS disease spread, and the role of calcium rather than iron accumulation in predicting future demyelination. We also highlight a potentially important role of low-myelin borders, that are known to connect to multiple areas within the M1 architecture, in disease spread. Importantly, the distinct pathology profile of a very-slow progressing patient (P4) highlights a distinction between disease duration and pathology progression. Our findings demonstrate the importance of *in-vivo* histology for the diagnosis and prognosis of neurodegenerative diseases such as ALS.

## Introduction

Amyotrophic lateral sclerosis (ALS) is a rapidly progressing neurodegenerative disease resulting in the loss of motor control, with a median survival time of 3 years^1,2^. It affects both upper (UMN) and lower (LMN) motor neurons^1^, where symptoms initially present focally in one body part. The disease spreads topographically, often first to the contralateral limb^3^, eventually resulting in bulbar symptoms which are associated with poorer outcomes^4^. Current knowledge of ALS pathology is largely based on *post-mortem* evidence^5–7^, therefore reflecting advanced disease stages. A detailed understanding of early-stage ALS pathology is needed to understand disease mechanisms and facilitate earlier diagnosis and prognosis. Such advances may identify novel therapeutic targets to slow disease progression.

*Post-mortem* studies have revealed abnormalities in the primary motor cortex (M1) in ALS, such as the depopulation of Betz cells in cortical layer 5(b)^5^, iron accumulation in deep cortex^6^ and increased intracellular calcium^7^. Since standard 1.5T or 3T MRI can hardly detect these layer-specific features, our understanding of *in-vivo* ALS pathology is limited and disease mechanisms are still debated, such as the focal hit versus multifocal pathology hypotheses^8^. Recent advances in ultra-high field MRI at 7T or above enable the automated assessment of anatomically-relevant cortical layers *in-vivo*^9–11^, allowing us to achieve *in-vivo* histology. The present study aims to employ this approach to provide a systematic description of *in-vivo* M1 pathology in early-stage ALS.

Previous MRI studies have shown iron accumulation in M1 of ALS-patients^12–14^, which has been approximately localised to deep M1, where it reflects neuroinflammation and microglia activation^6,15^, in addition to UMN impairment^16^. Moreover, iron accumulation strongly affects the topographic area corresponding to the symptom onset site (*i.e.,* first-affected)^17–19^. We previously showed that the topographic areas of M1 are microstructurally-distinct in healthy adults and can therefore be referred to as cortical fields^10^, emphasising the need for precise localisation. Moreover, we showed that the lower-limb (LL), upper-limb (UL) and face (F) fields show distinct levels of age-related iron accumulation^10^, highlighting the importance of using age-matched controls. In addition to iron, calcium is also dysregulated in ALS, where increased calcium is associated with glutamate excitotoxicity and motor neuron death in animal models^20–21^. This excitotoxicity may reflect the increased excitability of human M1 in ALS^22^. It is currently unclear how calcium dysregulation relates to iron accumulation and demyelination in ALS.

The low-myelin borders that separate cortical fields in M1^23–24^ show interruptions in myelination at the depths where the Betz cells are located^10,23^, which are known to be affected in ALS^5^, and are associated with different functional networks compared to cortical fields^26^. Since maladaptive proteins often propagate along highly-myelinated neurons in neurodegeneration^25^, these low-myelin borders may act as natural boundaries to limit disease spread. Alternatively, the degeneration of these borders may contribute to topographic disease spread between cortical fields^24^. An investigation into the stability of these low-myelin borders in ALS is therefore of interest.

The present study aimed to provide a systematic description of *in-vivo* pathology in ALS-patients. We used topographic layer imaging^24^ to characterise pathology with respect to both cortical layers and fields^10^. Sub-millimeter qT1 and QSM data, along with functional localisers, were collected from 12 ALS-patients and 12 age-, gender-, education- and handedness-matched controls. We used increased positive QSM (pQSM) values as a marker of iron^6,15^, increased qT1 values as a marker of (de)myelination^27^, and decreased (more negative) negative QSM (nQSM) as a marker of calcium^28^. We extracted microstructural profiles from each cortical layer (superficial, layer 5a, layer 5b, layer 6) according to a previously published approach^9–10^, and calculated multimodal *in-vivo* pathology maps. We then tested whether (i) the mean cortical thickness is decreased and/or the layer architecture of M1 is degenerated in patients compared to controls, (ii) iron accumulation is layer-specific in ALS, (iii) the accumulation of iron and calcium is topographic (*i.e.,* largely restricted to the first-affected cortical field) or atopographic (*i.e.,* not restricted to the first-affected cortical field), (iv) iron and/or calcium accumulation hotspots precede demyelination, (v) the low-myelin borders remain intact or disrupt over disease progression and show specifically high or specifically low substance accumulation and (vi) a very-slow progressing patient shows a distinct pathology profile compared to other patients.

## Materials and methods

### Participants

12 ALS-patients (P, 6 females, age: *M* = 60.5, *SD* = 12.7) and 12 healthy controls (C, 6 females, age: *M* = 61.1, *SD* = 11.9) took part in the present study between June 2018 and December 2022. Controls were individually matched to patients based on age (± 2 years; *t*(22) = −.15, *p* = .883), handedness, gender and years of education (± 4 years; patients: *M* = 14.5, *SD* = 2.7; controls: *M* = 15.4, *SD* = 2.7; *t*(22) = −.82, *p* = .422). On average, the revised ALS Functional Rating Scale (ALSFRS-R) score was 39.1 (*SD =* 6.5, range: 25-47) and patients were measured within 3 months of diagnosis (*SD* = 5, range: 0-14) and within 19 months of symptom onset (*SD* = 13, range: 2-48). Note that the latter excludes one patient (P4) with a very long disease duration (166 months between diagnosis and MRI). Participants underwent 7T-MRI and behavioural assessments. 3 patients (P1, P2, and P4) were measured again within 1 year (T2), while 1 patient (P4) was also measured 8 months after T2 (T3). Patients were recruited from the University Clinic Magdeburg, Hannover Medical School and Rostock University Medical Center (see **Table 1**). An experienced neurologist performed clinical assessments (see **Table 2**). Of the 12 patients, 7 had upper limb-onset (UL-onset; 3 right-lateralized, 2 left-lateralized, 1 bilateral), 2 had lower limb-onset (LL-onset; both left-lateralized) and 3 had bulbar-onset (B-onset; considered bilateral). Healthy controls were recruited from the DZNE database in Magdeburg, and exclusion criteria included sensorimotor deficits, neurologic disorders and 7T-MRI contraindications. All participants gave written informed consent and were paid. The study was approved by the local Ethics Committee of the Medical Faculty of the University of Magdeburg.

**Table 1.** Clinical and demographic information for ALS-patients (n = 12). Gender: F = female, M = male; Handedness: R = right, L = left; Onset type: UL = upper limb, LL = lower limb, B = bulbar; Onset side: L = left, R = right, B = bilateral; Phenotype: LMND = lower motor neuron dominant; UMN = upper motor neuron dominant, classical = both upper and lower motor neuron impairment. The Penn Upper Motor Neuron Scale (PUMNS)^32^ score indicates clinical signs of upper motor neuron involvement, with higher scores indicating greater impairment (M = 6.92, SD = 7.46, range: 0-22). The El-Escorial criteria indicates the degree of certainty of ALS diagnosis^28^. MRI-Symptoms indicates the time (in months) between the MRI and symptom onset, while MRI-Diagnosis indicates the time (in months) between MRI and formal diagnosis. Note that a negative MRI-Diagnosis indicates that the patient was scanned prior to a formal diagnosis. * Note that P4 first presented with an UL-isolated LMND phenotype, before progressing to a classical phenotype after 3 years from symptom onset.

| Patient | Age (years) | Education (years) | Gender | Handedness | Onset type | Onset side | Phenotype | PUMNS | El-Escorial | MRI-Symptoms (months) | MRI-Diagnosis (months) |
| --- | --- | --- | --- | --- | --- | --- | --- | --- | --- | --- | --- |
| P1 | 50 | 13 | F | R | UL | L | LMND | 1 | Possible | 12 | 5 |
| P2 | 66 | 18 | M | R | UL | L | UMND | 3 | Probable | 12 | 0 |
| P3 | 60 | 12 | M | L | UL | L | UMND | 11 | Definite | 12 | 0 |
| P4 | 48 | 13 | M | R | UL | R | Classical* | 2 | Definite | 189 | 166 |
| P5 | 77 | 14 | F | R | UL | R | LMND | 0 | Definite | 29 | 14 |
| P6 | 73 | 15 | M | R | UL | B | Classical | 9 | Definite | 14 | 5 |
| P7 | 74 | 15 | F | R | LL | L | UMND | 22 | Definite | 17 | 0 |
| P8 | 36 | 16 | M | R | LL | L | Classical | 15 | Probable | 28 | 4 |
| P9 | 52 | 12 | F | R | UL | R | UMND | 16 | Possible | 48 | 0 |
| P10 | 53 | 21 | M | R | B | B | Classical | 2 | Definite | 32 | 12 |
| P11 | 64 | 13 | F | R | B | B | LMND | 1 | Possible | 7 | 0 |
| P12 | 72 | 12 | F | R | B | B | LMND | 1 | Definite | 2 | -5 |

**Table 2.** Clinical assessment scores for ALS-patients (*n* = 12). ALSFRS-R = ALS Functional Rating Scale - Revised; KC stage = King’s College Staging; CNS-LS = Center for Neurologic Study-Lability Scale; CNS-BFS = Center for Neurologic Study - Bulbar Function Scale; Disease Progression Rate = 48-(ALSFRS-R total)/MRI-Symptoms. The ALSFRS-R^29^ indicates disease severity, where lower values indicate greater impairment, with subscores for fine, gross and bulbar motor function. The King’s College (KC) Stage^58^ indicates the stage of disease progression based on the ALSFRS-R score^59^, where stage 2A reflects the involvement of one body part and that a clinical diagnosis has taken place, while stage 2B and stage 3 reflect the subsequent involvement of second and third body parts, respectively. The CNS-LS^30^ indicates the frequency of pseudobulbar episodes, with higher scores indicating greater impairment. The CNS-BFS^31^ indicates bulbar dysfunction, with higher scores indicating greater impairment.

| Patient | ALSFRS-R total | ALSFRS-R fine motor | ALSFRS-R gross motor | ALSFRS-R bulbar | KC stage | CNS-LS | CNS-BFS | DPR |
| --- | --- | --- | --- | --- | --- | --- | --- | --- |
| P1 | 44 | 9 | 11 | 12 | 2A | 14 | 22 | 0.24 |
| P2 | 41 | 9 | 11 | 9 | 2B | 11 | 31 | 0.64 |
| P3 | 40 | 7 | 10 | 12 | 2A | 15 | 32 | 0.73 |
| P4 | 37 | 6 | 7 | 12 | 2B | 7 | 21 | 0.06 |
| P5 | 35 | 2 | 9 | 12 | 2A | 7 | 24 | 0.48 |
| P6 | 34 | 2 | 8 | 12 | 2B | 7 | 23 | 1.00 |
| P7 | 25 | 6 | 3 | 7 | 3 | 30 | 53 | 1.77 |
| P8 | 45 | 12 | 9 | 12 | 2A | 7 | 22 | 0.12 |
| P9 | 42 | 10 | 9 | 12 | 2B | 19 | 22 | 0.13 |
| P10 | 33 | 9 | 9 | 7 | 3 | 10 | 62 | 0.45 |
| P11 | 47 | 12 | 12 | 11 | 2A | 25 | 28 | 0.17 |
| P12 | 46 | 12 | 12 | 10 | 2A | 7 | 32 | 1.00 |

### 7T-MRI Data Acquisition

Data were collected using a 7T-MRI scanner (MAGNETOM, Siemens Healthcare) equipped with a 32-Channel Nova Medical Head Coil, located in Magdeburg, Germany. We acquired whole-brain, 0.7 mm isotropic resolution MP2RAGE images^34^ (sagittal slices, repetition time = 4800 ms, echo time = 2.01 ms, field-of view read = 224 mm, GRAPPA 2, flip angle = 5◦/3◦, invtime TI1/TI2 = 900/2750 ms, bandwidth = 250 Hz/Px). We also acquired whole-brain, 0.5 mm isotropic resolution susceptibility-weighted (SWI) images in 10/12 patients (transversal slices, repetition time = 22 ms, echo time = 9 ms, field-of view read = 192 mm, GRAPPA 2, flip angle = 10◦, bandwidth = 160 Hz/Px), since 2 patients could not be scanned any longer due to fatigue and discomfort (note that the SWI sequence was measured last). We also acquired a whole-brain, 1.5 mm isotropic resolution functional image (81 slices, repetition time = 2000 ms, echo time = 25 ms, field-of-view read = 212 mm, GRAPPA 2, interleaved acquisition) using an EPI gradient-echo BOLD sequence. The functional imaging involved a blocked-design paradigm with 12-second periods of body part movements (left/right foot, left/right hand, tongue) alternated with 15-seconds of rest (as described previously^10^). Before scanning, participants were trained in the movements and patients with difficulty were encouraged to conduct a reduced amount of movement precisely. Instructions inside the scanner (gray background, black color) asked participants to prepare (*e.g.,* ‘prepare right foot’) before performing the movement (*e.g.,* ‘move right foot’). Each movement was repeated four times, resulting in a total of 20 trials. Participants and patients wore fingerless braces covering the hands and forearms to reduce large movements of the hands and to facilitate movements of the patients. The total scanning duration was approximately 75 minutes.

### Image Processing

#### Structural Preprocessing

CBS Tools (v3.0.8)^35^ in MIPAV (v7.3.0)^36^ was used to process the structural data. Skull stripping and dura estimation were used to remove extra-cranial tissue, which was refined using ITK-SNAP (v3.8.0). The TOADS algorithm^37^ was used to segment the brain into different tissue types, before the CRUISE module was used to estimate tissue boundaries^38^, resulting in level set images. The distance field module was used to calculate mean cortical thickness (*i.e.*, across cortical depth). The volumetric layering module^39–40^ was used to divide the cortex into 21 depths, according to the equivolume approach. M1 masks were manually delineated using ITK-SNAP, as described previously^10^.

#### Quantitative Susceptibility Mapping (QSM)

After data quality checks, the SWI data of 4 patients (and the 4 matched controls) were excluded due to severe motion and truncation artefacts, leaving 16 participants for the SWI analyses. QSMbox (v2.0) was used to reconstruct QSM from the SWI images^41^. In line with previous studies^12,42^, QSM values were not normalised. We divided the QSM data into positive QSM (pQSM) and negative QSM (nQSM) values (as previously described^42^).

### Defining Cortical Layers

Using the simple curvature function in ParaView (v5.8.0), we calculated the mean curvature of the cortex. We then performed a vertex-wise linear regression analysis to predict qT1 values from mean curvature for each cortical layer in M1, on a subject specific basis, according to an existing approach^43^. The raw residuals of the regression models are referred to as ‘decurved’ qT1, following our previously published approach^10^. We then applied a data-driven approach to identify cortical layers in M1 from group-averaged decurved qT1 profiles separately for patients and controls^9–10^ (see **Fig.1B** for details, see **Supplementary Fig.1** for right hemisphere).

**Figure 1.**
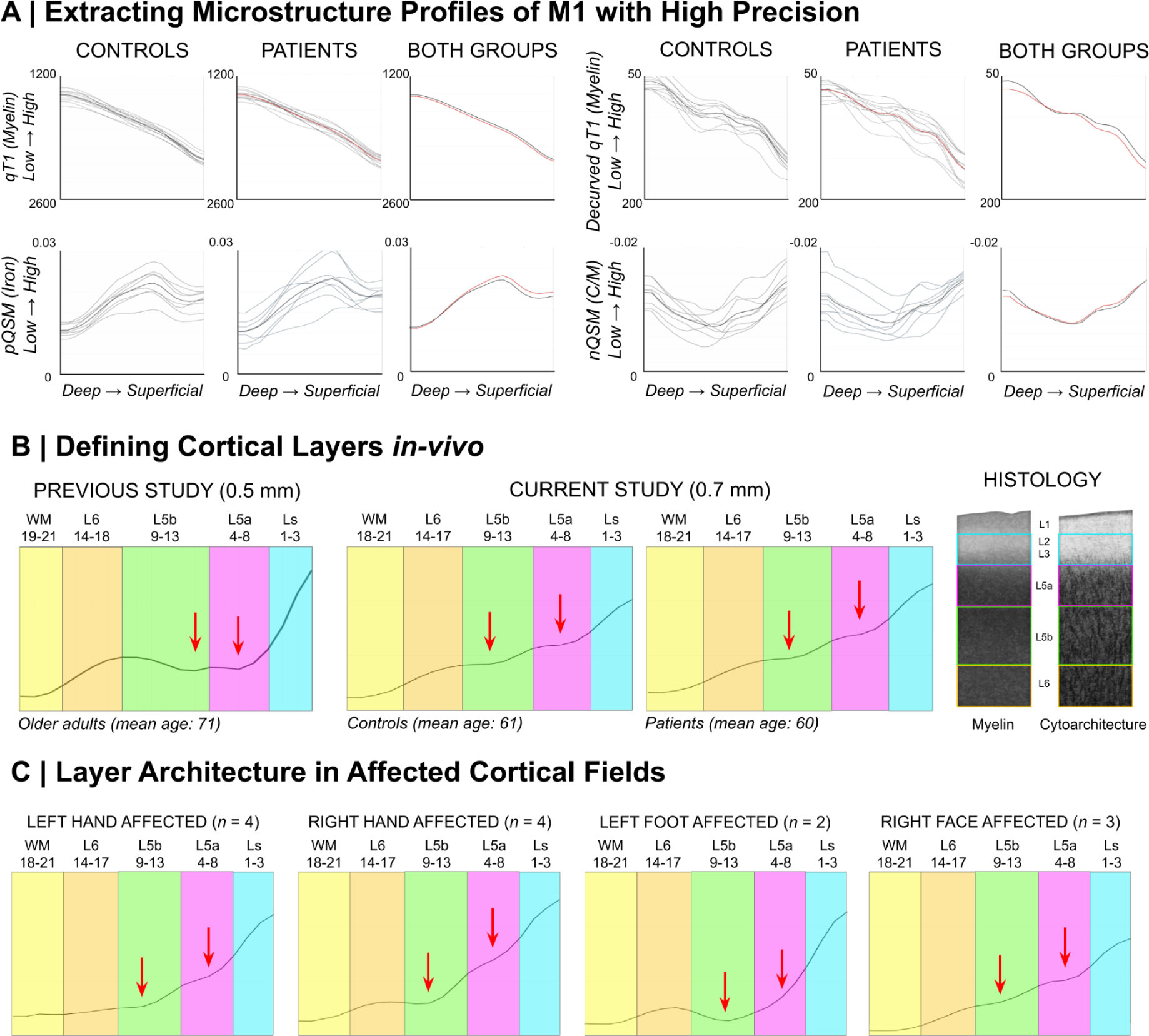
Microstructure profiles of the left primary motor cortex (M1) in ALS-patients and matched controls. **(A)** ‘Raw qT1’ (*i.e.,* not decurved; *n* = 12), decurved qT1 (*n* = 12), positive QSM (pQSM; *n* = 8) and negative QSM (nQSM; *n* =8) data extracted across all cortical depths (*n* = 21) of left M1. The first and second columns show data for all controls and all patients, with the group mean plotted in bold black and red, respectively. The third column shows the mean group data of the controls and patients, with red lines representing the patients. Note that qT1 and pQSM are validated *in-vivo* markers of myelin^27^ and iron^6,15^ content, respectively, while nQSM is largely considered to reflect calcium^28^ content. Also note that the profiles here represent the data of the entire M1, while in other figures and statistics the data used were averaged across cortical fields. **(B)** According to a previously published approach^9^, we identified four compartments (‘layers’) based on the ‘decurved qT1’ profile: Ls - superficial layer including layers 2-3; L5a - layer 5a; L5b - layer 5b; L6 - layer 6. Note that Ls does not include layer 1 since it is inaccessible with MRI^9^ or layer 4 since it is absent in M1. We show the layer definitions for healthy controls (*n* = 12) and ALS-patients (*n* = 12) in the present study (*centre*), as well as for older adults (*n* = 18) in our previous study^10^ (*left*). L5a and L5b were distinguished based on the presence of two small qT1 dips at the plateau of ‘decurved qT1’ values (indicating L5), while L6 was identified based on a sharp decrease in values before a further plateau indicating the presence of white matter. We show our layer approximations over schematic depictions (*right*) of M1 myelin^55^ and cell histological staining^56^. **(C)** Despite differences in the shape of the profile in the affected fields compared to the average across fields in patients, layers can be similarly identified in the affected fields.

### Functional Data Processing

The functional data were motion-corrected after acquisition using the Siemens ‘MoCo’ correction, before preprocessing in Statistical Parametric Mapping 12 (SPM12) which included smoothing (2 mm FWHM) and slice-timing correction. Coregistration was performed using the automated registration tool in ITK-SNAP, with further manual refinement based on anatomical landmarks where necessary. A first-level analysis created t-statistic maps (t-maps) for each body part (*e.g.,* left hand) based on contrast estimates (*e.g.,* [1 0 0 0 0]). To create the subject-specific functional localisers of the cortical fields (*e.g.,* left hand), we took the peak cluster of each t-map and removed overlapping voxels between localisers, as previously described^10^. The functional localisers were then mapped onto the same subject-specific cortical surfaces as used for the structural data. We extracted layer-wise qT1 and signed QSM values from the cortical fields, resulting in microstructural profiles for each cortical layer (4), each cortical field (3) and each hemisphere (2).

### Myelin Border Analysis

We identified myelin borders between the UL and F representations based on the highest qT1 value (*i.e.,* lowest myelin) located between the peak t-values of the UL and F localisers, using a previous approach^23,44^. For each cortical layer, we extracted qT1 and signed QSM values from the myelin borders and calculated the average qT1 and signed QSM values in the UL and F body part representations (UL+F/2), in order to compare values in the myelin border to the cortical fields.

### Layer-Specific Multimodal *in-vivo* Pathology Mapping

Using topographic layer imaging, we created multimodal *in-vivo* pathology maps on inflated cortical surfaces for each individual patient, in reference to the respective matched control (see **Fig.3** and **Supplementary Fig.2**). Pathology maps were calculated to identify disease hotspots indicated by demyelination (increased qT1, +1*SD* to +4*SD*s), iron accumulation (increased pQSM, +1*SD* to +4*SD*s) and calcium accumulation (more negative nQSM, −1*SD* to −4*SD*s) in each patient, relative to the mean M1 value of the matched control. The reference values were layer-specific to control for microstructural differences between layers in M1^10^. In addition, pathology maps are shown with individually-localised cortical fields in M1, to account for microstructural differences between cortical fields^10^.

### Behavioural Tests of Motor Function

All participants underwent body part-specific behavioural tests of motor function (as previously described^10^). To quantify lower limb (LL) function, the 6-Minute-Walking-Test (6MWT) was used to measure walking distance^45^. To quantify upper limb (UL) function, a dynamometer^46^ was used to measure hand strength, and the Purdue^47^ (Lafayette Instrument, model 32020A), Grooved^48^ (Lafayette Instrument, model 32025) and O’Connor^49^ (Lafayette Instrument, model 32021) pegboards were used to measure hand dexterity. To quantify bulbar (face, F) function, we used an automated tool that extracts features (*e.g.,* errors) of lateral tongue movements from short video clips^50^.

### Statistical Analyses

Statistical analyses were performed using IBM SPSS Statistics (v26, IBM, USA). Given the characteristic impaired motor function in ALS-patients, we used one-tailed paired-samples t-tests to test for group differences in motor behaviour. Based on evidence of demyelination, iron and calcium accumulation in ALS^5–7^, we used uncorrected one-tailed paired-samples t-tests to test for differences in cortical microstructure (*i.e.,* qT1, QSM) between groups. Uncorrected p-values were used to avoid false-negatives due to the small sample size. We also report effect sizes with 95% confidence intervals. Note that paired-samples t-tests were used to account for dependencies in the data (*i.e.,* matched patient-control pairs). We also used one-sample t-tests to test for differences in cortical microstructure between a single, particularly slow progressing patient (P4) and all other patients. We used linear regression models to test whether the layer-specific qT1, pQSM or nQSM predict whether or not a given cortical field is behaviourally affected. Individualised pathology maps were calculated by thresholding surfaces to show demyelination (increased qT1, +1*SD* to +4*SD*s), iron accumulation (increased pQSM, +1*SD* to +4*SD*s) and calcium accumulation (more negative nQSM, −1*SD* to −4*SD*s) in each patient relative to the mean M1 value of the matched control. Pathology estimations based on 1-4 *SD* are often used in clinical diagnostic imaging tools as they provide a detailed overview of how much the respective pathology differs from a control brain (*e.g.,* AIRAmed Software, https://www.airamed.de/de/startseite).

## Data availability

The data used in the present study have been made available in a public repository (https://github.com/alicianorthall/In-vivo-Pathology-ALS).

## Results

### Impaired Motor Function in ALS-patients

We quantified body part-specific motor function in 12 ALS-patients and 12 matched controls (LL: 6MWT; UL: hand strength, pegboards; F: tongue kinematics). For 3 patients, follow-up behaviour measurements were performed. The results show that ALS-patients are significantly impaired in motor function compared to matched controls (see **Supplementary Table 1**).

#### No Significant M1 Differences in Patients When Averaged Across Depths

We first tested whether the averaged qT1 and QSM values (averaged across cortical fields (LL, UL, F) and layers (L6, L5b, L5a, Ls) of M1 were significantly different in ALS-patients compared to controls to align with previous analyses^12,16^. Using this approach, we show no significant differences in pQSM values between patients and controls in left M1 (patients: *M* = .0172, *SD* = .0026; controls: *M* = .0171, *SD* = .0028; *t*(7) = 0.2, *p* = .428, *d* = −0.037, 95% CI [−1.02 0.94]) or right M1 (patients: *M* = .0185, *SD* = .0027; controls: *M* = .0172, *SD* = .0029; *t*(7) = 1.3, *p* = .114, *d* = −0.46, 95% CI [−1.46 0.53]). There are also no significant differences in nQSM between patients and controls in left M1 (patients: *M* = −.0094, *SD* = .0019; controls: *M* = −.0085, *SD* = .0013; *t*(7) = 1.6, *p* = .081, *d* = 0.55, 95% CI [−.45 1.55]) or right M1 (patients: *M* = −.0089, *SD* = .0021; controls: *M* = −.0087, *SD* = .0018; (*t*(7) = 0.3, *p* = .375, *d* = 0.10, 95% CI [−.88 1.08]). There are also no significant differences in qT1 between patients and controls in left M1 (patients: *M* = 1763.04 *SD* = 82.86; controls: *M* = 1783.64, *SD* = 50.62; (*t*(11) = 0.7, *p* = .764, *d* = .30, 95% CI [−.54 1.14]) or right M1 (patients: *M* = 1786.33, *SD* = 103.15; controls: *M* = 1816.21, *SD* = 49.35; *t*(11) = 1.0, *p* = .834, *d* = 0.37, 95% CI [−.47 1.21]). Finally, there are no significant differences in mean cortical thickness between patients and controls, although there is a trend towards significantly reduced cortical thickness in the patients compared to controls in the F field (see **Supplementary Table 2**).

### Layer-Specific M1 Pathology in ALS-patients

#### M1 Layer Architecture in ALS-patients

As in our previous work^10^, we used decurved qT1 values to identify four layers in M1 (Ls = superficial layer; L5a = layer 5a; L5b = layer 5b; L6 = layer 6; see **Fig.1B** for details). Our data show that those 4 layers can be identified similarly for ALS-patients and controls, where layers showed the same peaks (*e.g.,* level set 11 for L5b) and relative thicknesses (*e.g.,* level set 9-13 for L5b). In addition to the thickness, the principle layer architecture of M1 therefore also appears to be preserved in patients with early-stage ALS (see **Fig.1B**), also in the affected cortical fields (see **Fig. 1C**) although there are differences in the profile shape.

#### Pathological Substance Accumulation in M1 is Layer-Specific

We extracted qT1 and QSM values with high precision (see **Fig.1A**). Iron accumulation is a hallmark feature of ALS pathology^6,15^ and has been shown to particularly affect the deep region of M1. We show a significant increase in pQSM, specifically in L6 of left and right M1, in patients compared to matched controls (left M1: patients: *M* = .0144, *SD* = .0020; controls: *M* = .0128, *SD* = .0027; *t*(7) = 2.00, *p* = .043, *d* = 0.67, 95% CI [−0.33 1.68]; right M1: patients: *M* = .0165, *SD* = .0025; controls: *M* = .0140, *SD* = .0019; *t*(7) = 2.00, *p* = .041, *d* = 1.13, 95% CI [0.07 2.18]). These effects are moderate to large in size (see **Fig.2**). We show no significant differences or trends towards significance in pQSM between groups in the other layers (L5b, L5a and Ls; see **Table 3** for statistics).

**Figure 2.**
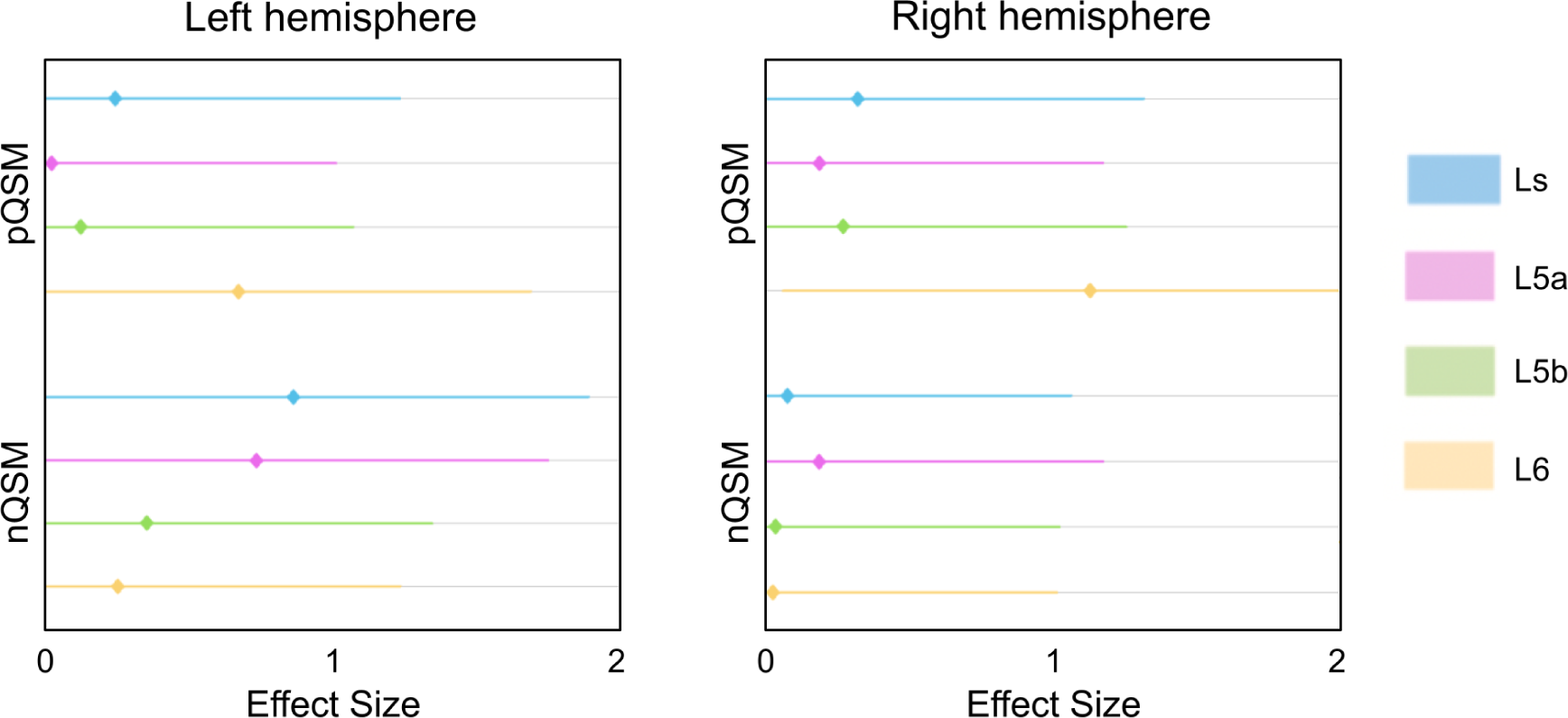
Layer-specific differences in QSM between ALS-patients (*n* = 8) and matched healthy controls (*n* = 8). One-tailed paired-samples t-tests were used to investigate matched patient-control pair differences in microstructure (pQSM, nQSM, qT1) for each cortical layer (see **Table 3** for statistics). Positive effect sizes indicate ‘more substance’ in patients compared to controls. Effect sizes (Cohen’s *d*) greater than 0.5 indicate medium effects, while effect sizes greater than 0.8 indicate large effects^60^. L6 - layer 6, L5b - layer 5b, L5a - layer 5a, Ls - superficial layer.

**Table 3.**
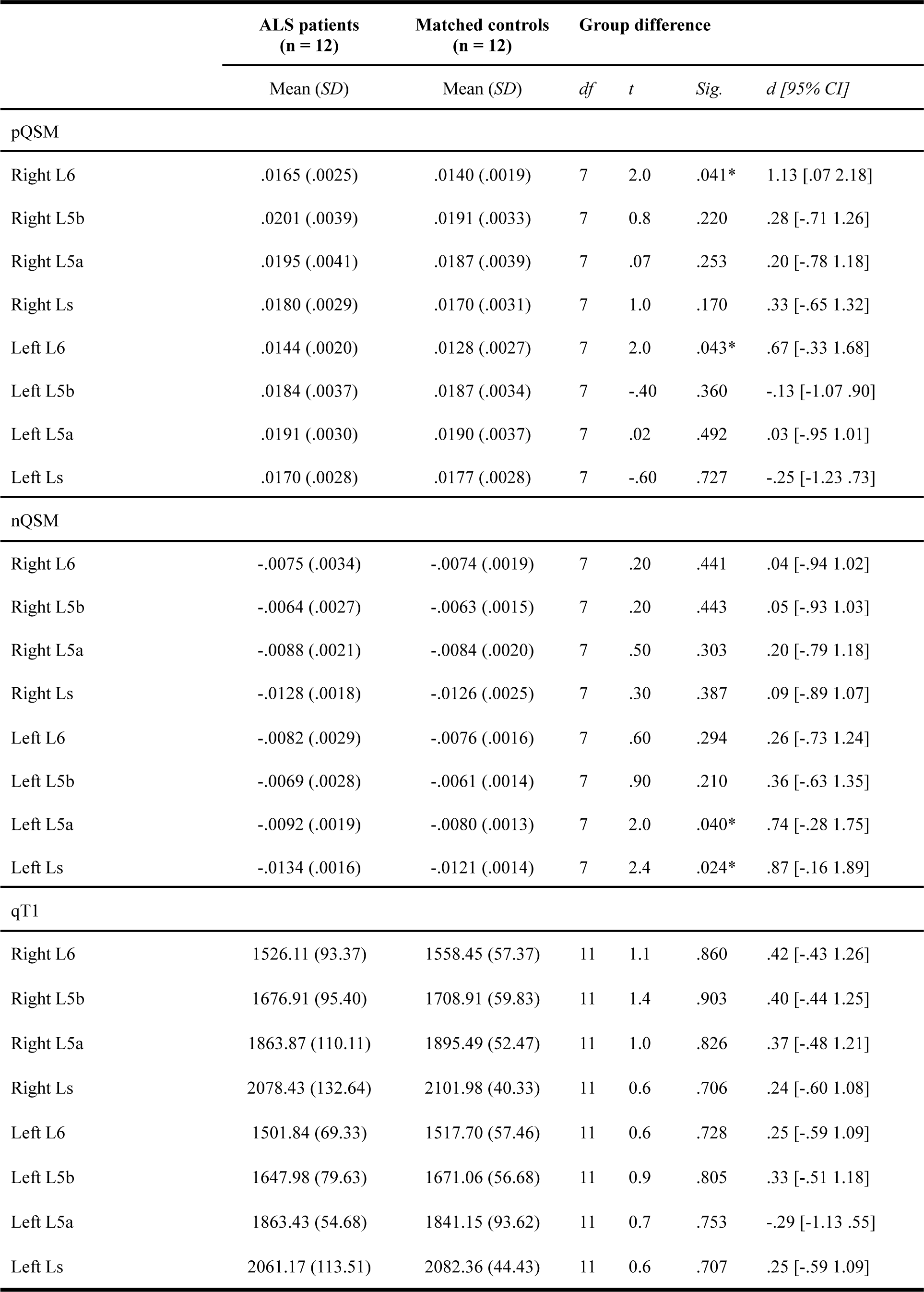
Layer-specific differences in M1 microstructure between patients (qT1: *n* = 12, QSM: *n* = 8) and matched healthy controls (qT1: *n* = 12, QSM: *n* = 8). One-tailed paired-samples t-tests were used to investigate matched patient-control pair differences in microstructure (pQSM, nQSM, qT1) for each cortical layer. * indicates significance at the .05 uncorrected level, while no tests show significance at the Bonferroni-corrected level (.05/32 = .002). Note that effect sizes (Cohen’s *d*) greater than 0.5 indicate medium effects, while effect sizes greater than 0.8 indicate large effects^60^. L6 - layer 6, L5b - layer 5b, L5a - layer 5a, Ls - superficial layer.

#### Iron Accumulates in First-Affected Cortical Field

Using topographic layer imaging, we created *in-vivo* pathology maps for each individual ALS-patient in reference to his/her matched control (see **Fig.3**, see **Supplementary Fig.2** for patients with qT1 data only). These maps reveal that demyelination is minimal compared to iron and calcium accumulation. As expected^17–19^, iron accumulation is higher in the first-affected cortical field compared to the other fields (excluding the contralateral field given evidence of highly symmetric pQSM increases^19^; see filled red arrows in **Fig.3**). This effect is specific to L6 (first-affected: *M* = 8.95, *SD* = 7.94, others: *M* = 2.49, *SD* = 1.68; *t*(6)= 2.81, *p* = .013, *d* = .95, 95% CI [−.04 2.22]), while there are no significant differences in L5b (first-affected: *M* = 6.66, *SD* = 8.19, others: *M* = 2.78, *SD* = 2.60; *t*(6)= 1.82, *p* = .056, *d* = .54, 95% CI [−.27 1.50]), L5a (first-affected: *M* = 4.66, *SD* = 3.56, others: *M* = 3.54, *SD* = 2.45; *t*(6)= 1.50, *p* = .089, *d* = .31, 95% CI [−.25 .95]) or Ls (first-affected: *M* = 3.11, *SD* = 2.44, others: *M* = 3.02, *SD* = 1.94; *t*(6)= .17, *p* = .436, *d* = .03, 95% CI [−.57 .65]). However iron accumulation also occurs in unaffected body parts (non-filled red arrows in **Fig.3**). An exception is P12, who does not show iron accumulation, but shows demyelination in the F field corresponding to the bulbar-onset type. Linear regression analyses show that pQSM pathology (nor qT1 pathology or nQSM pathology) does not significantly predict whether or not a given cortical field is behaviourally affected (see **Supplementary Table 3**).

**Figure 3.**
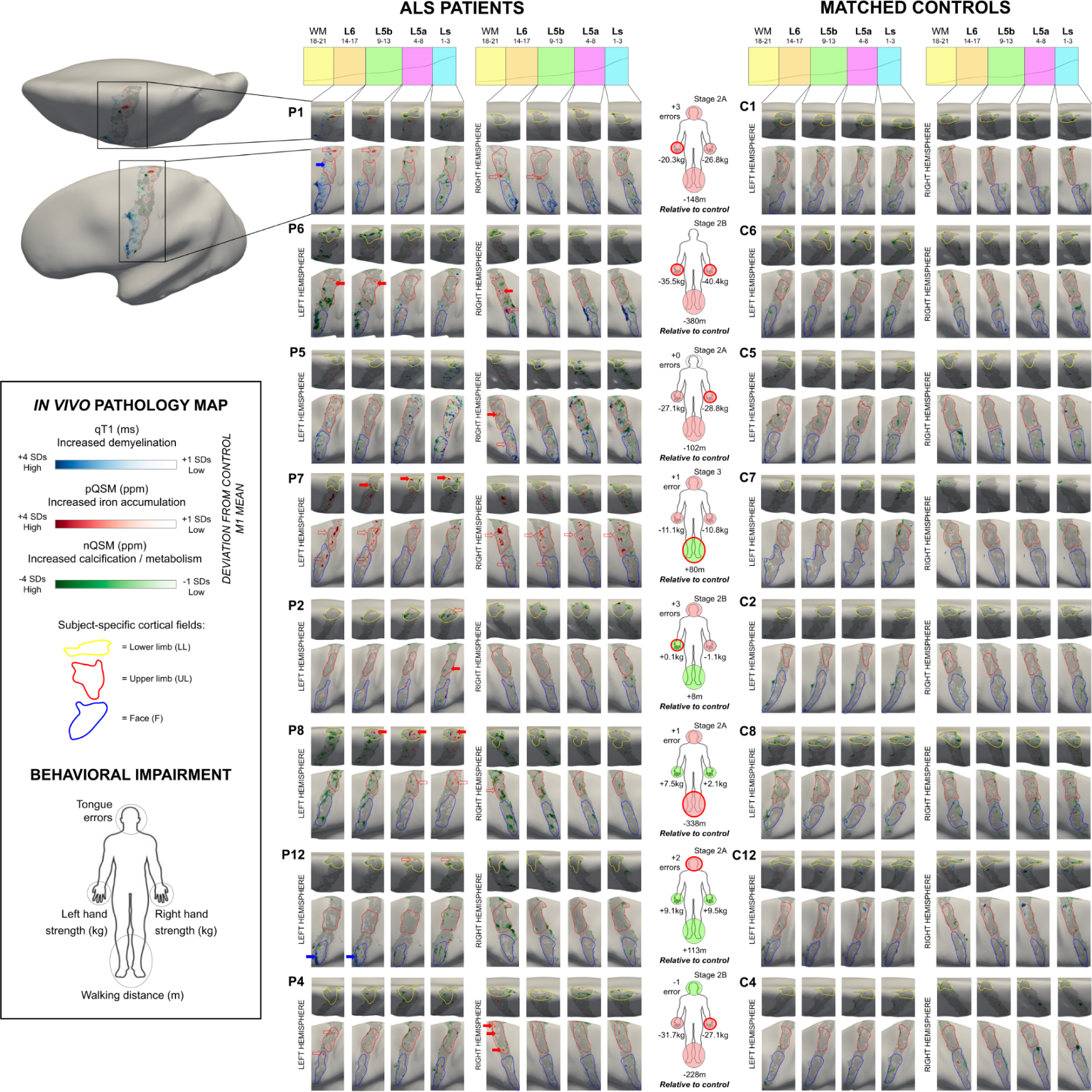
Multimodal *in-vivo* pathology maps in ALS-patients. Pathology maps were generated for each patient by thresholding the displayed value ranges at each layer to show increased qT1 and pQSM (+1 to +4 *SD*s), and reduced nQSM (−1 to −4 *SD*s), with respect to the mean M1 value of the matched control. Subject-specific cortical fields representing the lower limb, upper limb and face areas are outlined in yellow, red and blue, respectively. Pathology maps are shown for P1, P2, P8, P12, P6, P7, P4 and P5 (*left*) and their corresponding matched controls C1, C2, C8, C12, C6, C7, C4 and C5 (*right*). Pathology maps: note that filled red and blue arrows indicate iron accumulation and demyelination in the first-affected cortical field, respectively. Unfilled red arrows indicate iron accumulation in cortical fields other than the first-affected field. Body maps: note that red-outlined circles on the body maps indicate the onset site (*i.e.,* first-affected body part) of the patient. Filled red and green circles indicate impaired or better motor function in the circled body part compared to the matched control, respectively. The stage indicates the King’s College (KC) stage^58^ of disease progression based on ALSFRS-R score^59^: stage 2A reflects the involvement of one body part and that a clinical diagnosis has taken place, while stage 2B and stage 3 reflect the subsequent involvement of second and third body parts, respectively. L6 - layer 6, L5b - layer 5b, L5a - layer 5a, Ls - superficial layer.

#### Calcium Accumulation is Non-Topographic and Encompasses Low-Myelin Borders

The pathology maps (see **Fig.3**) provide an overview of calcium dysregulation in early-stage ALS, and suggest that calcium accumulation is atopographic. We show significantly decreased nQSM in the low-myelin borders (between the UL and F fields) compared to the cortical fields themselves (averaged UL and F fields), specifically in Ls (borders: *M* = −.0116, *SD* = .002; cortical fields: *M* = .0159, *SD* = .027; *t*(6) = −2.77; *p* = .033, *d* = −1.25, 95% CI [−2.68 .12]) and L5a (borders: *M* = −.0078, *SD* = .002; cortical fields: *M* = .0046, *SD* = .010; *t*(6) = −3.09; *p* = .021, *d* = −1.50, 95% CI [−3.11 .25]), but not in L5b (borders: *M* = −.0062, *SD* = .003; cortical fields: *M* = −.0015, *SD* = .018; *t*(6) = −.77; *p* = .468, *d* = −.30, 95% CI [−1.41 .72]) or L6 (borders: *M* = −.0074, *SD* = .004; cortical fields: *M* = −.0021, *SD* = .015; *t*(6) = −1.30; *p* = .242, *d* = −.42, 95% CI [−1.40 .45]).

#### Multiple Hotspots of Pathology in M1

We also addressed whether *in-vivo* pathology maps favour the hypothesis of one large pathology hotspot or multiple hotspots of pathology. The maps reveal that pathology occurs in multiple, small areas, often beyond the first-affected cortical field (see **Fig.3**). For example, P1 shows iron accumulation in an area between the left LL and UL fields compared to the control, while this patient shows demyelination in the (first-affected) left UL field.

#### Calcium Accumulation Precedes Demyelination

Out of the 12 patients, three (P1, P2 and P4) were also measured longitudinally (see **Fig.4**). To provide an overview over disease progression, we quantified the increase in the percentage of demyelinated vertices (+2*SD*s compared to matched control mean) in the first-affected cortical field between timepoints (see **Supplementary Table 4**). In addition, we show that the topography of pathological calcium (vertices with QSM values > than −2*SD*s from the matched control M1 mean) at T1 predicts myelination at T2 and T3, most strongly in Ls (see **Supplementary Table 5**). We also show that the topography of pathological iron (vertices with QSM values > +2*SD*s from the matched control M1 mean) at T1 predicts myelination at T2 (but not T3), specifically in L6 (see **Supplementary Table 6**). However, the latter effect did not survive correction for multiple comparisons.

**Figure 4.**
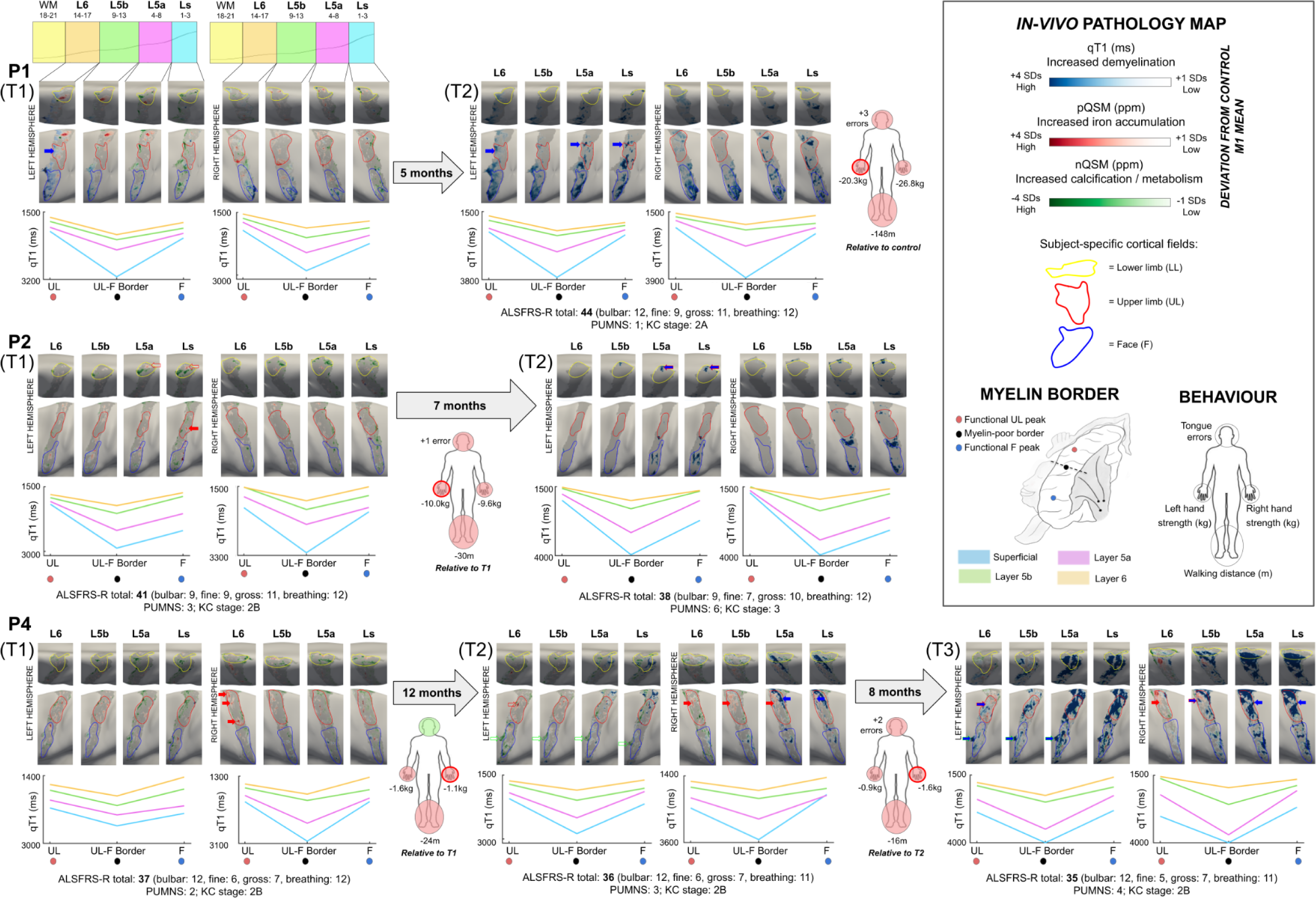
Longitudinal multimodal *in-vivo* pathology maps in ALS-patients. Individualised *in-vivo* pathology maps were generated by thresholding the displayed value ranges at each layer to show increased qT1 and pQSM (+1 to +4 *SD*s), and reduced nQSM (−1 to −4 *SD*s), with respect to the mean M1 value of the matched control at baseline. Row 1 shows the pathology maps for P1 at T1 (time point 1) and T2 (time point 2), row 2 shows the pathology maps for P2 at T1 and T2 and row 3 shows the pathology maps for P4 at T1, T2 and T3 (time point 3). Pathology maps: note that red and blue arrows on the pathology maps indicate iron accumulation and demyelination in the cortical field corresponding to symptom onset site, respectively. Body maps: note that red-outlined circles on the body maps indicate the onset site of the patient, while filled red and green circles indicate impaired or better motor function in the circled body part compared to the matched control, respectively. Note that QSM data were excluded at T2 for P1 and P2 due to severe artefacts. Clinical information: ALS Functional Rating Scale - Revised (ALSFRS-R)^29^ indicates disease severity, where lower values indicate greater impairment, with subscores for fine, gross and bulbar motor function. The Penn Upper Motor Neuron Scale (PUMNS)^32^ score indicates clinical signs of upper motor neuron involvement, with higher scores indicating greater impairment. The King’s College (KC) stage^58^ indicates disease progression based on ALSFRS-R score^59^: stage 2A reflects the involvement of one body part and that a clinical diagnosis has taken place, while stage 2B and stage 3 reflect the subsequent involvement of second and third body parts, respectively. L6 - layer 6, L5b - layer 5b, L5a - layer 5a, Ls - superficial layer.

#### Disrupted Low-Myelin Borders With Disease Progression

We also addressed whether the low-myelin borders that separate the F and UL cortical fields in M1 are perhaps particularly affected or rather preserved in ALS to identify their role in disease progression. The longitudinal pathology maps show demyelination of the F-UL border at T2 compared to T1 (see P1, P2 and P4 in **Fig.4**), and at T3 compared to T2 (see P4 in **Fig.4**). Based on visual inspection and the quantified demyelination (see **Supplementary Table 4**), this effect is largely restricted to Ls and L5a, where the low-myelin borders are typically most defined in M1 of healthy adults^10,23^.

#### Multimodal MRI Markers of Slow Disease Progression

To identify cortical markers related to slow disease progression, we compared the individualized *in-vivo* pathology maps between the very slow-progressing patient (P4) and all other patients. We show that P4 shows a significantly higher percentage of abnormal pQSM vertices (*i.e.,* increased iron accumulation) in L5b (*t*(6) = 17.07, *p* = 10^−5^, *d* = 6.45, 95% CI [2.84 10.08]) and L5a (*t*(6) = 5.53, *p* = .001, *d* = 2.09, 95% CI [.70 3.43]), in addition to a significantly higher percentage of abnormal nQSM vertices (*i.e.,* increased calcium accumulation) in L5a (*t*(6) = 4.96, *p* = .003, *d* = 1.88, 95% CI [.58 3.13]) at T1, compared to all other patients. In addition, our data show that P4 shows a significantly lower percentage of abnormal qT1 vertices (*i.e.,* less demyelination) in all layers compared to the other patients at T1 (L6: *t*(10) = −2.40, *p* = .037, *d* = −.73, 95% CI [−1.38 −.04]; L5b: *t*(10) = −2.79, *p* = .019, *d* = −.75, 95% CI [−1.41 −.06]; L5a: *t*(10) = −2.33, *p* = .042, *d* = −.70, 95% CI [−1.35 −.02]; Ls: *t*(10) = −2.40, *p* = .038, *d* = −.72, 95% CI [−1.38 −.04]).

## Discussion

ALS is a rapidly-progressing neurodegenerative disease characterised by the loss of motor control^1^. The present study aimed to systematically describe the *in-vivo* pathology of M1 in early-stage ALS-patients. Given the layer-specific and topographic signature of pathology in ALS^5^, we combined submillimetre structural 7T-MRI data (qT1, QSM) and automated layer modelling with cortical field-specific analyses to calculate precise *in-vivo* histology profiles of the patients. Our data reveal critical insights into the *in-vivo* pathology of ALS that can be summarised as (i) preserved mean cortical thickness and preserved layer architecture of M1, (ii) layer-specific iron (layer 6) and calcium (layer 5a, superficial layer) accumulation, (iii) topographic iron accumulation, but atopographic calcium accumulation, (iv) disrupted low-myelin borders with increased calcium, (v) later demyelination may occur more in the earlier hotspots of calcium accumulation, and does therefore not show a strictly topographic profile, (v) demyelination characterises disease progression (particularly in layer 5a and superficial layer), and (vi) the very slow-progressing patient has a distinct pathology profile in M1 compared to the other patients. Overall, our study provides novel insights into the *in-vivo* M1 pathology in ALS, offering new perspectives into disease mechanisms and diagnosis (see **Fig.5**).

**Figure 5.**
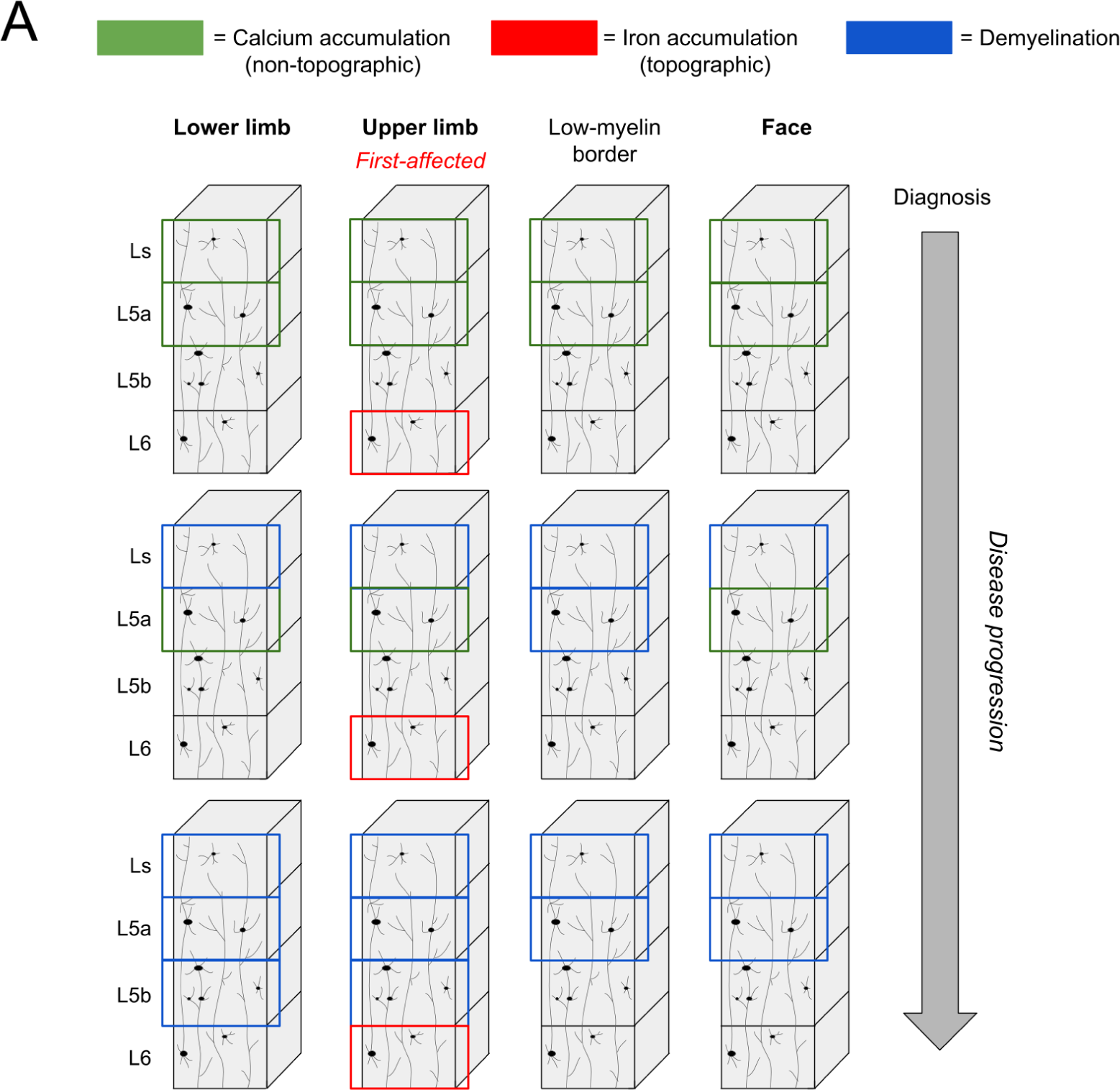
Model of *in-vivo* M1 pathology progression in ALS. Schematic depiction of the layer-specific pathology features shown in ALS patients compared to matched healthy controls in the present study. In the early pathology stage, we demonstrate topographic (*i.e.,* most in first-affected cortical field - example upper limb shown here) iron accumulation in L6 and non-topographic (*i.e.,* more widespread) calcium accumulation in Ls and L5a (see Table 3, see Fig.2 and Fig.3). With disease progression, we highlight increasing demyelination (see Supplementary Table 4), particularly in Ls and L5a and in the low-myelin borders between adjacent cortical fields (see Fig.4, based on visual inspection), corresponding to calcium accumulation at earlier time points (see Supplementary Table 5). L6 - layer 6, L5b - layer 5b, L5a - layer 5a, Ls - superficial layer.

Across cortical depth in M1, we show no significant differences in microstructure or mean cortical thickness between patients and controls. This contrasts previous evidence of reduced cortical thickness in ALS-patients compared to healthy controls at varying MR field strengths^16,61–63^. However, the patients in the present study show a relatively slower disease progression rate compared to a previous 7T study^16^. Moreover, our cortical thickness estimates of healthy controls are comparable to previous automated 7T estimates^16,64^. We show a trend towards reduced mean cortical thickness in the F field of patients compared to controls. This field-specific trend may reflect a higher vulnerability of this field to neurodegeneration, since it has been shown that myelination is reduced in the F field compared to the other cortical fields of M1^10^. Our findings therefore confirm that layer-specific microstructural pathology in ALS is detectable in the absence of significant cortical atrophy.

Layer-specific analyses reveal that iron accumulation in ALS is specific to L6 of M1. Although this effect was moderate to large in size, it is critical to replicate this finding in a larger cohort of patients given the high heterogeneity of the disorder. This extends previous studies showing deep iron accumulation in ALS, which is considered to reflect neuroinflammation and activated microglia^6,15^. This suggests that iron accumulation in L6 may provide an earlier marker of pathology than the degeneration of L5(b)^5^. Our data also show that iron accumulates most strongly in the first-affected cortical field, as expected^17–19^, and that this is specific to L6. However, in our data, the degree of iron accumulation (% of pQSM pathology) does not predict whether or not a given cortical field is behaviourally affected. This may suggest that iron accumulation largely serves as a specific marker for the first-affected cortical field, which may support early diagnosis, and that this marker is a consequence rather than the cause of the disease.

Surface mapping reveals multiple, small hotspots of pathology rather than one large hotspot. For example, one patient (P1) shows L6 iron accumulation in an area between the LL and UL fields, which may reflect pathology in the torso region where impairment is difficult to detect. Another patient (P2) shows iron accumulation in Ls-L5a of the first-affected (UL) field, deviating from the expected location in L6, but in all layers of the contralateral F field. This may reflect the quantitatively impaired tongue function in this patient, highlighting challenges in detecting bulbar symptoms clinically^50^. This demonstrates potential mismatches between brain and behaviour in ALS^53^, therefore emphasising the utility of MRI for individualised medicine.

We also investigated calcium dysregulation, demonstrating that Ls (including L2-L3) and L5a show calcium accumulation in ALS. In addition, we show that calcium accumulation is higher in the areas between cortical fields, where the low-myelin borders are located, compared to the cortical fields themselves. This highlights the atopographic profile of calcium accumulation in ALS, in contrast to the more topographic iron accumulation. Previous evidence has shown disrupted calcium homeostasis in ALS^7^, where increased intracellular calcium is associated with glutamate excitotoxicity in animal models^20–21^. Increased calcium in humans may reflect similar mechanisms, including the increased activity of M1 in ALS^51^ or calcifications in blood vessels related to poor brain health^52^. Interestingly, patients with pronounced calcium accumulation (P8, P12) largely show more focal behavioural impairment in our data. This may suggest that calcium accumulation provides an early disease marker which may be related to compensation, as has been suggested for increased activity in M1^51^.

Based on a small sample of patients with longitudinal data, we provide evidence towards widespread demyelination of M1 with disease progression. We also show that later demyelination is higher at previous calcium accumulation hotspots, whereas the relation to previous iron accumulation hotspots is weaker. Moreover, the low-myelin borders between cortical fields that are disrupted with disease progression show earlier increased calcium accumulation compared to cortical fields. Overall, this points towards a partly atopographic character of disease spread in ALS, and a potential role of the low-myelin borders, which are connected to multiple body parts^23,26^, in disease spread. These findings also suggest that metabolism disruption and inflammation precede cell loss in ALS^54^. . An alternative perspective to this interpretation is that connections between low-myelin borders and subcortical structures^26^ are more affected by pathology, compared to connections between cortical fields and subcortical structures. From this perspective, the affected border areas would not reflect markers of disease spread, but would be the consequence of affected subcortical systems. Further investigation is needed to characterise the role of these atopographic disease mechanisms in ALS.

Finally, we also investigated whether the in-vivo pathology profile of the very-slow progressing patient (P4) is distinct from the other patients. We demonstrate that P4 shows increased iron accumulation in L5b and L5a, in addition to increased calcium accumulation in L5a, compared to other patients. Moreover, P4 shows decreased demyelination in all layers compared to the other patients. These results suggest that P4, despite the long disease duration, may be in the earlier stage of cortical pathology (see **Fig.5**) that is characterised by increased substance accumulation. One may interpret this finding to argue that there are distinct pathology profiles for disease duration (*i.e.,* longer survival associated with substance accumulation) and disease progression (*i.e.,* increased disease severity associated with demyelination). More specifically, one may argue that the “turning point” between substance accumulation and cell loss is delayed in the slow-progressing patient, rather than a delay in substance accumulation as such. These assumptions have to be verified in a larger patient sample with more slow- and more fast-progressing patients to avoid overinterpretation based on single patient profiles.

With respect to the applied methodology, although the method we used to define cortical layers in M1 is based on an *in-vivo-ex-vivo* validation model^9–10^, we cannot guarantee that our layer definitions correspond to the exact biological layers. This limitation should be addressed with more extensive *in-vivo-ex-vivo* validation studies. Finally, although quantitative and validated markers of cortical microstructure were used here, we cannot be certain of the exact tissue properties measured.

In summary, we provide novel insights into the *in-vivo* pathology of ALS-patients using topographic layer imaging. We show that layer-specific changes in tissue microstructure precede gross cortical atrophy. We highlight the topographic nature of iron accumulation as a marker for diagnosis, while atopographic calcium accumulation may precede demyelination and cell loss. The role of the low-myelin borders between cortical fields, which show particularly high calcium accumulation, in disease progression needs further clarification. Finally, we highlight the distinct pathology profile of a very-slow progressing patient, where increased substance accumulation and reduced demyelination may indicate a lack of pathology progression despite a long disease duration. Our efficient 7T-MRI scanning protocol, and use of open-source analysis tools^35^, make our approach accessible to large-cohort investigations.

## Acknowledgements

The authors would like to thank the volunteer patients, and their families, for their participation in the study.

## Author Contributions

E.K. and S.S. designed the study. S.S, S.P. and J.P. recruited the patients. A.N., M.W. and J.D. collected the data. S.V. performed the clinical assessments. J.D., A.N. and I.T. developed the processing pipelines. A.N. completed the formal analysis, data investigation and statistical analysis. A.N. wrote the manuscript. E.K., J.D., S.S. and M.W. reviewed and edited the manuscript. E.K. and S.S. supervised the study. All authors contributed to the article and approved the submitted version.

## References

1. Al-Chalabi A, Hardiman O, Kiernan MC, Chiò A, Rix-Brooks B, van den Berg LH. (2016). Amyotrophic lateral sclerosis: moving towards a new classification system. The Lancet Neurology. 2016;15(11),1182-1194.

2. Norris, F., Shepherd, R., Denys, E., Kwei, U., Mukai, E., Elias, L.,…&Norris, H. (1993). Onset, natural history and outcome in idiopathic adult motor neuron disease. Journal of the neurological sciences, 118(1), 48–55.

3. Ravits JM, La Spada AR. ALS motor phenotype heterogeneity, focality, and spread: deconstructing motor neuron degeneration. Neurology. 2009;73(10),805–811.

4. Chio A, Logroscino G, Hardiman O, et al. Prognostic factors in ALS: a critical review. Amyotrophic lateral sclerosis. 2009;10(5-6),310–323.

5. Hammer RP, Tomiyasu U, Scheibel AB. Degeneration of the human Betz cell due to amyotrophic lateral sclerosis. Experimental neurology. 1979;63(2),336–346.

6. Kwan JY, Jeong SY, Van Gelderen P, et al. Iron accumulation in deep cortical layers accounts for MRI signal abnormalities in ALS: correlating 7 tesla MRI and pathology. PloS one. 2012;7(4),e35241.

7. Appel SH, Beers D, Siklos L, Engelhardt JI, Mosier DR. Calcium: the darth vader of ALS. Amyotrophic Lateral Sclerosis and Other Motor Neuron Disorders. 2001;2(1),47–54.

8. Sekiguchi T, Kanouchi T, Shibuya K, et al. Spreading of amyotrophic lateral sclerosis lesions—multifocal hits and local propagation?. Journal of Neurology, Neurosurgery & Psychiatry. 2014;85(1),85–91.

9. Huber L, Handwerker DA, Jangraw DC, et al. High-resolution CBV-fMRI allows mapping of laminar activity and connectivity of cortical input and output in human M1. Neuron. 2017;96(6),1253–1263.

10. Northall A, Doehler J, Weber M, Vielhaber S, Schreiber S, Kuehn E. Layer-specific vulnerability is a mechanism of topographic map aging. Neurobiology of Aging. 2023;128,17–32.

11. Doehler J, Northall A, Liu P, Fracasso A, Chrysidou A, Speck O, Lohmann G, Wolbers T, Kuehn E. The 3D Structural Architecture of the Human Hand Area Is Nontopographic. Journal of Neuroscience. 2023;43(19),3456–3476.

12. Acosta-Cabronero J, Machts J, Schreiber S, et al. Quantitative susceptibility MRI to detect brain iron in amyotrophic lateral sclerosis. Radiology. 2018a;289(1),195–203.

13. Adachi Y, Sato N, Saito Y, et al. Usefulness of SWI for the detection of iron in the motor cortex in amyotrophic lateral sclerosis. Journal of Neuroimaging. 2015;25(3),443–451.

14. Wang C, Foxley S, Ansorge O, et al. Methods for quantitative susceptibility and R2* mapping in whole post-mortem brains at 7T applied to amyotrophic lateral sclerosis. NeuroImage. 2020;222,117216.

15. Pallebage-Gamarallage M, Foxley S, Menke RA, et al. Dissecting the pathobiology of altered MRI signal in amyotrophic lateral sclerosis: A post mortem whole brain sampling strategy for the integration of ultra-high-field MRI and quantitative neuropathology. BMC neuroscience. 2018;19(1),1–24.

16. Cosottini M, Donatelli G, Costagli M, et al. High-resolution 7T MR imaging of the motor cortex in amyotrophic lateral sclerosis. American journal of neuroradiology. 2016;37(3),455–461.

17. Costagli M, Donatelli G, Biagi L, et al. Magnetic susceptibility in the deep layers of the primary motor cortex in amyotrophic lateral sclerosis. NeuroImage: Clinical. 2016;12,965–969.

18. Donatelli G, Ienco EC, Costagli M, et al. MRI cortical feature of bulbar impairment in patients with amyotrophic lateral sclerosis. NeuroImage: Clinical. 2019;24,101934.

19. Donatelli G, Costagli M, Cecchi P, et al. Motor cortical patterns of upper motor neuron pathology in amyotrophic lateral sclerosis: A 3 T MRI study with iron-sensitive sequences. NeuroImage: Clinical. 2022;35,103138.

20. Kruman II, Pedersen WA, Springer JE, Mattson MP. ALS-linked Cu/Zn–SOD mutation increases vulnerability of motor neurons to excitotoxicity by a mechanism involving increased oxidative stress and perturbed calcium homeostasis. Experimental neurology. 1999;160(1),28–39.

21. Siklós L, Engelhardt J, Harati Y, Smith RG., Joó F, Appel SH. Ultrastructural evidence for altered calcium in motor nerve terminals in amyotrophic lateral sclerosis. Annals of neurology. 1996;39(2),203–216.

22. Eisen A, Pant B, Stewart H. Cortical excitability in amyotrophic lateral sclerosis: a clue to pathogenesis. Canadian journal of neurological sciences. 1993;20(1),11–16.

23. Kuehn E, Dinse J, Jakobsen E, et al. Body topography parcellates human sensory and motor cortex. Cerebral Cortex. 2017;27(7),3790–3805.

24. Schreiber S, Northall A, Weber M, Vielhaber S, Kuehn E. Topographical layer imaging as a tool to track neurodegenerative disease spread in M1. Nature Reviews Neuroscience. 2021;22(1),68–69.

25. Bartzokis G, Lu PH, Mintz J. Human brain myelination and amyloid beta deposition in Alzheimer’s disease. Alzheimer’s & dementia. 2007;3(2),122–125.

26. Gordon EM, Chauvin RJ, Van AN, et al. A somato-cognitive action network alternates with effector regions in motor cortex. Nature. 2023;1–9.

27. Stüber C, Morawski M, Schäfer A, et al. Myelin and iron concentration in the human brain: a quantitative study of MRI contrast. Neuroimage. 2014;93,95–106.

28. Wang Y, Spincemaille P, Liu Z, et al. Clinical quantitative susceptibility mapping (QSM): Biometal imaging and its emerging roles in patient care. Journal of magnetic resonance imaging. 2017;46(4),951–971.

29. Brooks BR, Miller RG, Swash M, Munsat TL. El Escorial revisited: revised criteria for the diagnosis of amyotrophic lateral sclerosis. Amyotrophic lateral sclerosis and other motor neuron disorders. 2000;1(5),293–299.

30. Cedarbaum JM, Stambler N, Malta E, et al. The ALSFRS-R: a revised ALS functional rating scale that incorporates assessments of respiratory function. Journal of the neurological sciences. 1999;169(1-2),13–21.

31. Moore SR, Gresham LS, Bromberg MB, Kasarkis EJ, Smith RA. A self report measure of affective lability. Journal of Neurology, Neurosurgery & Psychiatry. 1997;63(1),89–93.

32. Smith RA, Macklin EA, Myers KJ, et al. Assessment of bulbar function in amyotrophic lateral sclerosis: validation of a self-report scale (Center for Neurologic Study Bulbar Function Scale). European journal of neurology. 2018;25(7),907–e66.

33. Quinn C, Edmundson C, Dahodwala N, Elman L. Reliable and efficient scale to assess upper motor neuron disease burden in amyotrophic lateral sclerosis. Muscle & nerve. 2020;61(4),508–511.

34. Marques JP, Kober T, Krueger, G, van der Zwaag W, Van de Moortele PF, Gruetter R. MP2RAGE, a self bias-field corrected sequence for improved segmentation and T1-mapping at high field. Neuroimage. 2010;49(2),1271–1281.

35. Bazin PL, Weiss M, Dinse J, Schäfer A, Trampel R, Turner R. A computational framework for ultra-high resolution cortical segmentation at 7 Tesla. NeuroImage. 2014;93,201–209.

36. McAuliffe MJ, Lalonde FM, McGarry D, Gandler W, Csaky K, Trus BL. Medical image processing, analysis and visualization in clinical research. In Proceedings 14th IEEE Symposium on Computer-Based Medical Systems. 2001,381–386.

37. Bazin PL, Pham DL. Homeomorphic brain image segmentation with topological and statistical atlases. Medical image analysis. 2008;12(5),616–625.

38. Han X, Pham DL, Tosun D, Rettmann ME, Xu C, Prince JL. CRUISE: cortical reconstruction using implicit surface evolution. NeuroImage. 2004;23(3),997–1012.

39. Waehnert MD, Dinse J, Weiss M, et al. Anatomically motivated modeling of cortical laminae. Neuroimage. 2014;93,210–220.

40. Waehnert MD, Dinse J, Schäfer A, et al. A subject-specific framework for in vivo myeloarchitectonic analysis using high resolution quantitative MRI. Neuroimage. 2016;125,94–107.

41. Acosta-Cabronero J, Milovic C, Mattern H, Tejos C, Speck O, Callaghan MF. A robust multi-scale approach to quantitative susceptibility mapping. Neuroimage. 2018b;183,7–24.

42. Betts MJ, Acosta-Cabronero J, Cardenas-Blanco A, Nestor PJ, Düzel E. High-resolution characterisation of the aging brain using simultaneous quantitative susceptibility mapping (QSM) and R2* measurements at 7 T. Neuroimage. 2016;138,43–63.

43. Sereno MI, Lutti A, Weiskopf N, Dick F. Mapping the human cortical surface by combining quantitative T 1 with retinotopy. Cerebral cortex. 2013;23(9),2261–2268.

44. Glasser MF, Coalson TS, Robinson EC, et al. A multi-modal parcellation of human cerebral cortex. Nature. 2016;536(7615),171-178.

45. Enright PL. The six-minute walk test. Respiratory care. 2003;48(8),783–785.

46. Peolsson A, Hedlund R, Öberg B. Intra-and inter-tester reliability and reference values for hand strength. Journal of rehabilitation medicine. 2001;33(1),36–41.

47. Tiffin J, Asher EJ. The Purdue Pegboard: norms and studies of reliability and validity. Journal of applied psychology. 1948;32(3),234.

48. Matthews, C. G., & Klove, H. (1964). Instruction manual for the adult neuropsychology test battery. Madison, WI: University of Wisconsin Medical School, 36.

49. Fleishman EA. A modified administration procedure for the O’Connor Finger Dexterity Test. Journal of Applied Psychology. 1953;37(3),191.

50. Northall, A., Mukhopadhyay, B., Weber, M., Petri, S., Prudlo, J., Vielhaber, S.,…&Kuehn, E. (2022). An automated tongue tracker for quantifying bulbar function in ALS. Frontiers in Neurology, 223.

51. Proudfoot M, Bede P, Turner MR. Imaging cerebral activity in amyotrophic lateral sclerosis. Frontiers in neurology. 2019;9,1148.

52. Schreiber S, Bernal J, Arndt P, et al. Brain vascular health in ALS is mediated through motor cortex microvascular integrity. Cells. 2023;12(6),957.

53. Verstraete E, Turner MR, Grosskreutz J, et al. Mind the gap: the mismatch between clinical and imaging metrics in ALS. Amyotrophic lateral sclerosis and frontotemporal degeneration. 2015;16(7-8),524–529.

54. Toft MH, Gredal O, Pakkenberg B. The size distribution of neurons in the motor cortex in amyotrophic lateral sclerosis. Journal of anatomy. 2005;207(4),399–407.

55. Maranzano A, Verde F, Colombo E, Poletti B, Doretti A, Bonetti R, Gagliardi D, Meneri M, Corti S, Morelli C, Silani V & Ticozzi N. Regional spreading pattern is associated with clinical phenotype in amyotrophic lateral sclerosis. Brain. 2023; doi: 10.1093/brain/awad129.

56. Dinse J, Härtwich N, Waehnert MD, et al. A cytoarchitecture-driven myelin model reveals area-specific signatures in human primary and secondary areas using ultra-high resolution in-vivo brain MRI. Neuroimage. 2015;114,71–87.

57. Vogt C, Vogt O. Allgemeine Ergebnisse unserer Hirnforschung [General results of our brain research]. Zeitschrift für Augenheilkunde. 1919;25,273–462.

58. Roche JC, Rojas-Garcia R, Scott KM, Scotton W, Ellis CE, Burman R, Wijesekera L, Turner M, Leigh N, Shaw C, Al-Chalabi A. A proposed staging system for amyotrophic lateral sclerosis. Brain. 2012;135(3), 847–852.

59. Balendra R, Jones A, Jivraj N, Knights C, Ellis CM, Burman R, Turner M, Leigh N, Shaw C, Al-Chalabi A. Estimating clinical stage of amyotrophic lateral sclerosis from the ALS Functional Rating Scale. Amyotrophic Lateral Sclerosis and Frontotemporal Degeneration. 2014;15(3-4), 279–284.

60. Sawilowsky SS. New effect size rules of thumb. Journal of modern applied statistical methods. 2009;8(2), 26.

61. Agosta F, Valsasina P, Riva N, Copetti M, Messina MJ, Prelle A, Comi G, Filippi M. The cortical signature of amyotrophic lateral sclerosis. PloS one. 2012;16: 1–7.

62. Roccatagliata L, Bonzano L, Mancardi G, Canepa C, Caponnetto C. Detection of motor cortex thinning and corticospinal tract involvement by quantitative MRI in amyotrophic lateral sclerosis. Amyotrophic Lateral Sclerosis. 2009;10(1), 47–52.

63. Donatelli G, Retico A, Ienco EC, Cecchi P, Costagli M, Frosini D, Biagi L, Tosetti M, Siciliano G, Cosottini M. Semiautomated evaluation of the primary motor cortex in patients with amyotrophic lateral sclerosis at 3T. American Journal of Neuroradiology. 2018;39(1), 63–69.

64. Lüsebrink F, Wollrab A, Speck O. Cortical thickness determination of the human brain using high resolution 3 T and 7 T MRI data. Neuroimage. 2013;70, 122–131.

