## Supplementary materials for "Multimodal layer modelling reveals *in-vivo* pathology in ALS"

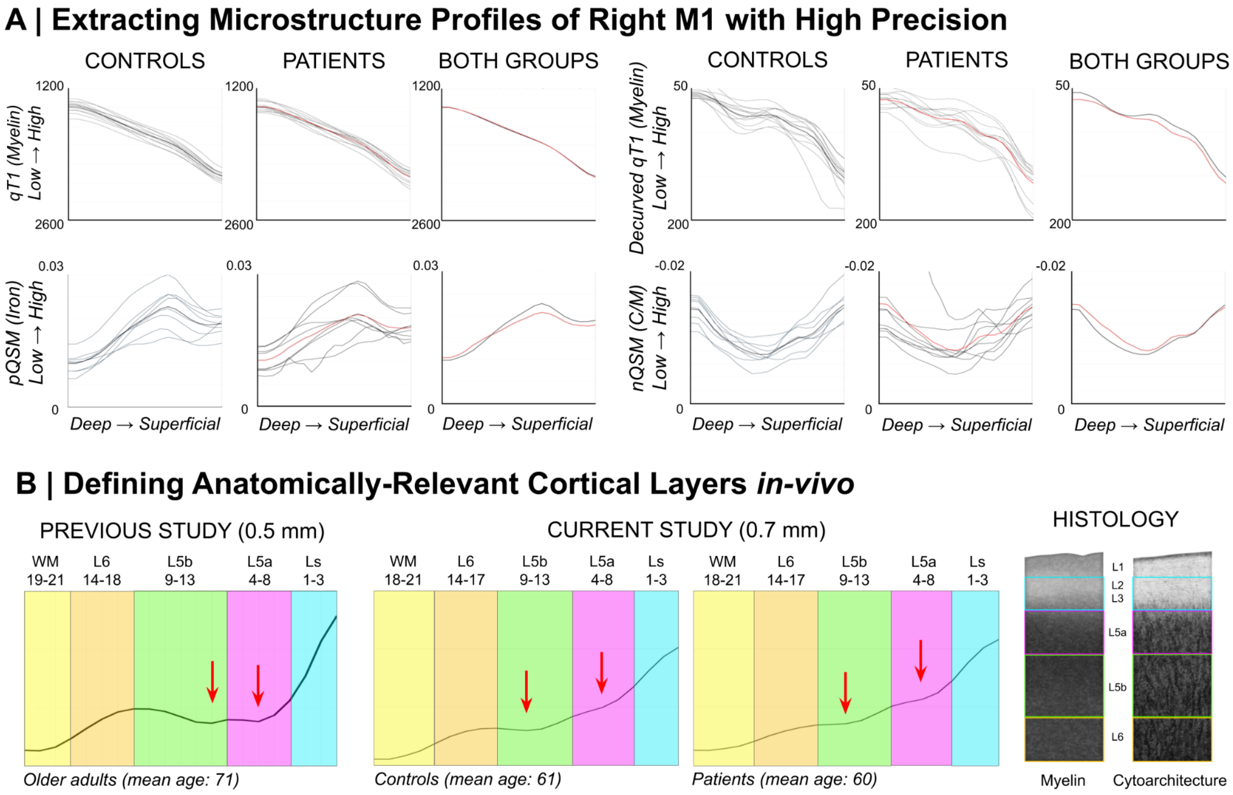


**Supplementary Figure 1. Microstructure profiles of the right primary motor cortex (M1) in ALS-patients and matched controls. (A)** 'Raw qT1' (*i.e.,* not decurved; *n* = 12), decurved qT1 (*n* = 12), positive QSM (pQSM; *n* =8) and negative QSM (nQSM; *n* = 8) data extracted across all cortical depths (*n* = 21) of left M1. The first two columns show data for all controls and all patients, with the group mean plotted in bold black and red, respectively. The third column shows the mean group data of the controls and patients, with red lines representing the patients. Note that qT1 and pQSM are validated *in-vivo* markers of myelin[27] and iron[6,15] content, respectively, while nQSM is largely considered to reflect calcium[28] content. Also note that the profiles here represent the data of the entire M1, while in other figures and statistics the data used were averaged across cortical fields. **(B)** According to a previously published approach[9], we identified four anatomically-relevant compartments ('layers') based on 'decurved qT1': Ls - superficial layer including layers 3 and 4; L5a - layer 5a; L5b - layer 5b; L6 - layer 6. We show these layer definitions for healthy controls (*n* = 12) and patients (*n* = 12) in the present study (*centre*), as well as for older adults (*n* = 18) in our previous study[10] (*left*). L5a and L5b were distinguished based on the presence of two small qT1 dips at the plateau of 'decurved qT1' values (indicating L5), while L6 was identified based on a sharp decrease in values before a further plateau indicating the presence of white matter. We show our layer approximations over schematic depictions (*right*) of M1 myelin[55] and cell histological staining[56] are shown.

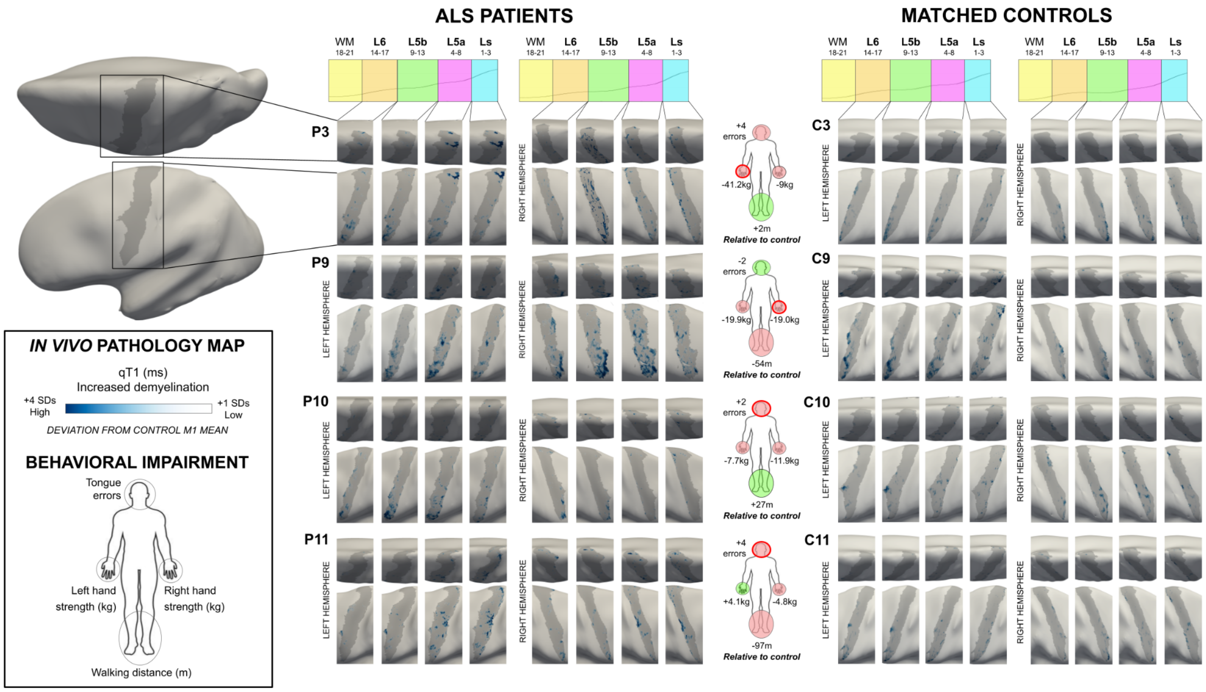


**Supplementary Figure 2. Individualised *in-vivo* demyelination maps in ALS-patients.** Demyelination maps were generated by thresholding the displayed qT1 value ranges at each layer to show increased qT1 (+1 to +4 SDs) with respect to the mean M1 value of the matched control. Maps are shown for P3, P9, P10 and P11, and their corresponding matched controls C3, C9, C10 and C11. Note that red-outlined circles on the body maps indicate the onset site of the patient, while filled red and green circles indicate impaired or better motor function in the circled body part compared to the matched control, respectively. L6 - layer 6, L5b - layer 5b, L5a - layer 5a, Ls - superficial layer.

**Supplementary Table 1.** Group differences in behavioural tests of motor function between ALS-patients (*n* = 12) and matched healthy controls (*n* = 12). One-tailed paired-samples t-tests were used to investigate group differences. * indicates significance at the .05 uncorrected level, while ** indicates significance at the Bonferroni-corrected level (.05/16 = .003).

| | **Patients** | **Controls** | **Group difference** | | | |
|---|---|---|---|---|---|---|
| | Mean (*SD*) | Mean (*SD*) | *df* | *t* | *Sig.* | *g [95% CI]* |
| 6MWT (m) | 446.03 (107.11) | 539.08 (73.72) | 11 | 2.05 | .032* | 1.02 [.16 1.86] |
| rHand strength (kg) | 19.72 (11.78) | 33.72 (8.82) | 11 | 3.30 | .004* | 1.35 [.46 2.23] |
| lHand strength (kg) | 17.16 (13.39) | 31.61 (9.53) | 11 | 2.88 | .007* | 1.24 [.37 2.12] |
| TT errors | 2.55 (1.81) | 1.00 (1.10) | 10 | -2.60 | .013* | -1.04 [-1.89 -.18] |
| rPurdue (sec) | 113.08 (90.20) | 61.38 (6.09) | 11 | -1.93 | .040* | -.81 [-1.64 .02] |
| rPurdue (drops) | 2.50 (3.66) | .50 (.67) | 11 | -1.81 | .049* | -.76 [-1.59 .07] |
| lPurdue (sec) | 147.58 (104.86) | 66.75 (8.90) | 11 | -2.70 | .010* | -1.09 [-1.94 -.23] |
| lPurdue (drops) | 2.92 (4.17) | .42 (.67) | 11 | -2.11 | .029* | -.84 [-1.67 $-10^{-3}$] |
| rGrooved (sec) | 122.00 (86.05) | 79.75 (11.26) | 11 | -1.64 | .065 | -.69 [-1.51 .14] |
| rGrooved (drops) | 2.00 (3.77) | .42 (.67) | 11 | -1.38 | .098 | -.59 [-1.40 .23] |
| lGrooved (sec) | 150.00 (100.05) | 80.75 (13.00) | 11 | -2.48 | .015* | -.97 [-1.82 -.13] |
| lGrooved (drops) | 2.42 (4.06) | 0.00 (0.00) | 11 | -2.06 | .032* | -.84 [-1.68 -.01] |
| rO'Connor (total) | 22.50 (12.60) | 32.25 (8.62) | 11 | 2.09 | .970 | .90 [.06 1.74] |
| rO'Connor (drops) | 12.75 (13.73) | 5.83 (3.46) | 11 | -1.68 | .060* | -.69 [-1.52 .13] |
| lO'Connor (total) | 21.83 (15.51) | 32.33 (7.83) | 11 | 2.47 | .985 | .86 [.02 1.69] |
| lO'Connor (drops) | 13.83 (14.94) | 4.25 (2.56) | 11 | -2.19 | .025* | -.89 [-1.73 -.06] |

**Supplementary Table 2.** Group differences in mean cortical thickness between ALS-patients and matched healthy controls. Mean cortical thickness (*i.e.,* across cortical depth) was calculated within each cortical field of the left hemisphere. One-tailed paired-samples t-tests were used to investigate group differences.

| | **ALS patients** | **Matched controls** | **Group difference** | | | |
|---|---|---|---|---|---|---|
| Cortical field | Mean (*SD*) | Mean (*SD*) | *df* | *t* | *Sig.* | *g [95% CI]* |
| Face (F) | 1.85 (.28) | 2.02 (.16) | 18 | -1.58 | .065 | -.71 [-1.65 .17] |
| Upper limb (UL) | 1.92 (.09) | 1.96 (.17) | 20 | -.48 | .318 | -.28 [-1.13 .55] |
| Lower limb (LL) | 1.97 (.09) | 1.97 (.18) | 20 | .07 | .471 | 0.00 [-.84 .84] |

**Supplementary Table 3.** Linear regression models were used to predict whether a given cortical field was behaviourally affected (affected = 1) or unaffected (affected = 0) from the percentage of qT1, pQSM or nQSM pathology (separate models for each). Note that a cortical field was marked 'affected' if one or more points were lost in the respective body part subscore of the ALSFRS-R (*i.e.,* fine motor subscore = UL; gross motor subscore = LL; bulbar subscore = F; see Table 2), according to the King's College (KC) staging criteria[58]. ALSFRS-R - ALS Functional Rating Scale - Revised; qT1 - quantitative T1; pQSM - positive QSM; nQSM - negative QSM; L6 - layer 6, L5b - layer 5b, L5a - layer 5a, Ls - superficial layer.

| **Cortical layer** | ***df*** | ***F*** | $R^2$ | ***p*** |
|---|---|---|---|---|
| qT1 pathology (T1) ~ affected/unaffected (T2) | | | | |
| L6 | 1, 68 | 2.73 | .04 | .103 |
| L5b | 1, 68 | 1.69 | .02 | .198 |
| L5a | 1, 68 | 2.33 | .03 | .132 |
| Ls | 1, 68 | 1.30 | .02 | .259 |
| pQSM pathology (T1) ~ affected/unaffected (T2) | | | | |
| L6 | 1, 46 | .48 | .01 | .492 |
| L5b | 1, 46 | 1.31 | .03 | .258 |
| L5a | 1, 46 | 2.02 | .04 | .162 |
| Ls | 1, 46 | 1.12 | .02 | .294 |
| nQSM pathology (T1) ~ affected/unaffected (T2) | | | | |
| L6 | 1, 46 | .04 | $10^{-5}$ | .842 |
| L5b | 1, 46 | .73 | .02 | .397 |
| L5a | 1, 46 | .50 | .01 | .484 |
| Ls | 1, 46 | $10^{-6}$ | $10^{-7}$ | .992 |

**Supplementary Table 4.** Difference in the percentage of demyelinated vertices (with qT1 values > + 2SDs from the matched control M1 mean) from T1 (time point 1) to T2 (timepoint 2) or T2 to T3 (time point 3). L6 - layer 6, L5b - layer 5b, L5a - layer 5a, Ls - superficial layer.

| | **Percentage change of demyelinated vertices in first-affected cortical field** | | | |
|---|---|---|---|---|
| | **L6** | **L5b** | **L5a** | **Ls** |
| P2<br>(T1 to T2) | .25% | .73% | 9.80% | 12.40% |
| P4<br>(T1 to T2) | .90% | .08% | 10.80% | 17.13% |
| P4<br>(T2 to T3) | 30.14% | 46.12% | 61.03% | 52.57% |

**Supplementary Table 5.** Linear regression models were used to predict follow-up (time point 2 (T2) or time point 3 (T3), separate models for each) qT1 values from pathological nQSM values (> -2SDs from matched control M1 mean; time point 1 (T1)). * indicates significance at the .05 uncorrected level, while ** indicates significance at the Bonferroni-corrected level (.05/12 = .004). L6 - layer 6, L5b - layer 5b, L5a - layer 5a, Ls - superficial layer.

| **Cortical layer** | ***df*** | ***F*** | $R^2$ | ***p*** |
|---|---|---|---|---|
| P4<br>nQSM (T1) ~ qT1 (T2) | | | | |
| L6 | 1, 58 | .64 | .003 | .425 |
| L5b | 1, 135 | .46 | .003 | .498 |
| L5a | 1, 201 | 1.21 | .006 | .272 |
| Ls | 1, 192 | 7.05 | .050 | .009* |
| P4<br>nQSM (T1) ~ qT1 (T3) | | | | |
| L6 | 1, 60 | .221 | .004 | .640 |
| L5b | 1, 135 | .376 | .003 | .541 |
| L5a | 1, 201 | .093 | $10^{-5}$ | .761 |
| Ls | 1, 192 | 8.66 | .043 | .003** |
| P2<br>nQSM (T1) ~ qT1 (T2) | | | | |
| L6 | 1, 10 | .018 | $10^{-5}$ | .895 |
| L5b | 1, 14 | 4.94 | .261 | .043* |
| L5a | 1, 112 | 6.22 | .053 | .014* |
| Ls | 1, 209 | 9.07 | .042 | .003** |

**Supplementary Table 6.** Linear regression models were used to predict follow-up (time point 2 (T2) or time point 3 (T3), separate models for each) qT1 values from pathological pQSM values (> +2SDs from matched control M1 mean; time point 1 (T1)). * indicates significance at the .05 uncorrected level, while ** indicates significance at the Bonferroni-corrected level (.05/12 = .004). L6 - layer 6, L5b - layer 5b, L5a - layer 5a, Ls - superficial layer.

| **Cortical layer** | ***df*** | ***F*** | $R^2$ | ***p*** |
|---|---|---|---|---|
| P4<br>pQSM (T1) ~ qT1 (T2) | | | | |
| L6 | 1, 175 | 5.98 | .033 | .016* |
| L5b | 1, 297 | .30 | .001 | .582 |
| L5a | 1, 426 | .87 | .002 | .352 |
| Ls | 1, 174 | 3.83 | .022 | .052 |
| P4<br>pQSM (T1) ~ qT1 (T3) | | | | |
| L6 | 1, 426 | 1.76 | .004 | .186 |
| L5b | 1, 297 | 0.59 | .002 | .444 |
| L5a | 1, 175 | 2.06 | .012 | .153 |
| Ls | 1, 174 | .35 | .002 | .553 |
| P2<br>pQSM (T1) ~ qT1 (T2) | | | | |
| L6 | 1, 106 | 8.23 | .072 | .005* |
| L5b | 1, 137 | .011 | $10^{-6}$ | .917 |
| L5a | 1, 184 | 2.71 | .015 | .101 |
| Ls | 1, 276 | 1.35 | .005 | .247 |
